# BioProfiling.jl: Profiling biological perturbations with high-content imaging in single cells and heterogeneous populations

**DOI:** 10.1101/2021.06.18.448961

**Authors:** Loan Vulliard, Joel Hancock, Anton Kamnev, Christopher W. Fell, Joana Ferreira da Silva, Joanna Loizou, Vanja Nagy, Loïc Dupré, Jörg Menche

## Abstract

**Motivation:** High-content imaging screens provide a cost-effective and scalable way to assess cell states across diverse experimental conditions. The analysis of the acquired microscopy images involves assembling and curating morphological measurements of individual cells into morphological profiles suitable for testing biological hypotheses. Despite being a critical step, there is currently no standard approach to morphological profiling and no solution is available for the high-performance Julia programming language.

**Results:** Here, we introduce BioProfiling.jl, an efficient end-to-end solution for compiling and filtering informative morphological profiles in Julia. The package contains all the necessary data structures to curate morphological measurements and helper functions to transform, normalize and visualize profiles. Robust statistical distances and permutation tests enable quantification of the significance of the observed changes despite the high fraction of outliers inherent to high-content screens. This package also simplifies visual artifact diagnostics, thus streamlining a bottleneck of morphological analyses. We showcase the features of the package by analyzing a chemical imaging screen, in which the morphological profiles prove to be informative about the compounds’ mechanisms of action and can be conveniently integrated with the network localization of molecular targets.

**Availability:** The Julia package is available on GitHub: https://github.com/menchelab/BioProfiling.jl We also provide Jupyter notebooks reproducing our analyses: https://github.com/menchelab/BioProfilingNotebooks

## Introduction

High-Content Screening (HCS) enables profiling cellular phenotypes across hundreds of thousands of conditions by combining automated microscopy with advanced image analysis methods. HCS thus represents a flexible and cost-effective solution for replacing multiple specific assays (Simm *et al*., 2018; Way *et al*., 2021; Chandrasekaran *et al*., 2020), and has been widely adopted in both basic and applied research. Notable achievements range from drug discovery (Simm *et al*., 2018; Chandrasekaran *et al*., 2020; Scheeder *et al*., 2018) to the elucidation of combinatorial drug effects (Caldera *et al*., 2019) and to ex-vivo drug-response screening in patients (Snijder *et al*., 2017). Depending on the application, the analysis of HCS experiments may involve a variety of tasks. For instance, one might perform a classification task to infer the mechanism of action of candidate drugs (Ljosa *et al*., 2013; Ando *et al*., 2017; Pawlowski *et al*., 2016), compare cellular phenotypes in various conditions (Gustafsdottir *et al*., 2013; German *et al*., 2020; Rohban *et al*., 2017) or describe interactions between cellular perturbations (Breinig *et al*., 2015; Caldera *et al*., 2019; Heigwer *et al*., 2018; Fischer *et al*., 2015; Billmann *et al*., 2016). All these cases involve numerous experimental and analytical steps.

A typical HCS experiment starts from preparing microplates with cells subjected to various perturbations, such as different drugs, and stained using standardized protocols, such as the Cell Painting assay (Bray *et al*., 2016) (Fig. 1A). These microplates are then imaged using automated confocal fluorescence microscopy, resulting in a large number of images. Each image is then analyzed to extract quantitative morphological measurements that describe the respective cellular phenotype. Some tools are commonly used for this numerical feature extraction step (Pau *et al*., 2010; McQuin *et al*., 2018), and recent deep learning approaches attempt to replace expert-curated measurements with data-driven discriminative features (Ando *et al*., 2017; Pawlowski *et al*., 2016; Lu *et al*., 2019).

**Fig. 1.**
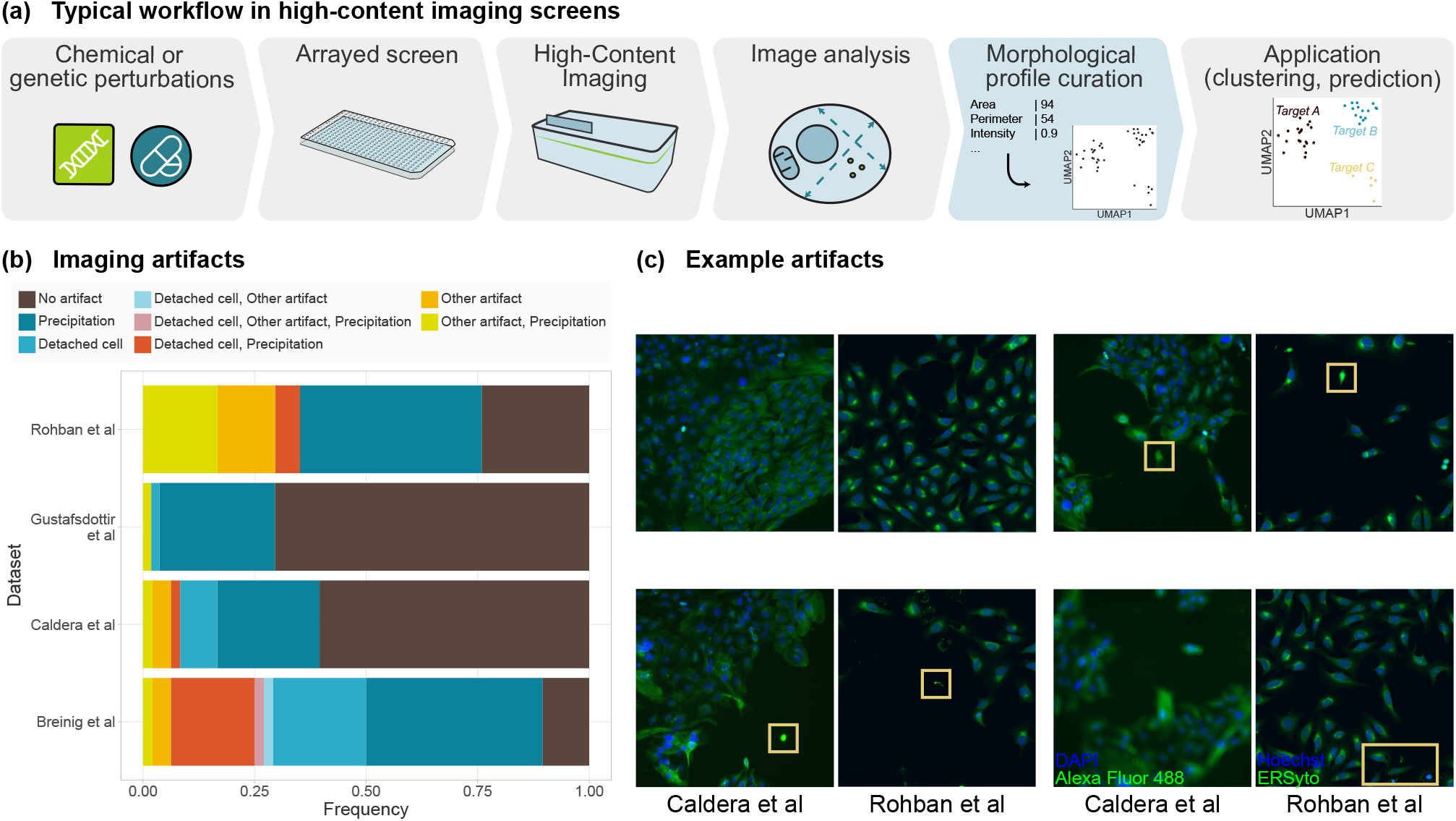
High-Content Screening (HCS) experiments require adequate analysis tools. **(a)** Standard analysis workflow of HCS experiments. **(b)** Quantification of imaging artifacts that may lead to biases in HCS analyses in sample images from four published studies (Rohban et al., 2017; Gustafsdottir et al., 2013; Caldera et al., 2019; Breinig et al., 2015). **(c)** Examples of such imaging artifacts. Boxes highlight regions of interest.

While the analytical tasks of HCS experiments vary between applications, they involve common data normalization and filtering steps, and guidelines have been proposed for computing informative representations of cellular phenotypes, usually referred to as morphological profiles (Caicedo *et al*., 2017; Bougen-Zhukov *et al*., 2017). A tool suitable to facilitate morphological profiling must meet several criteria. First, it must be versatile, to adapt to different HCS use cases and to cope with the diverse challenges inherent to such experiments (Boutros *et al*., 2015; Chandrasekaran *et al*., 2020; Ljosa *et al*., 2013; Caicedo *et al*., 2017). These challenges include technical problems such as blurred images, poorly adherent cells, saturated pixels, staining artifacts and segmentation mistakes. HCS studies need to address these frequent limitations, as in some experiments most images are affected (Figure 1b,c). Second, the analysis pipelines must account for background noise, intensity bias and potential confounders, including plate layout and batch effects. Third, the considerable heterogeneity of the morphological descriptors must be handled. Cellular morphology might vary greatly in the analysed cell populations due to the experimental setup, heterogeneous cell types or cell states, inconsistencies in perturbation efficiency, or when timing-dependent phenomena are imaged as snapshots.

The few actively-maintained tools attempting to fulfill these needs include CellProfiler Analyst and its graphical user interface, designed to handle CellProfiler measurements (McQuin *et al*., 2018; Jones *et al*., 2008), cellHTS2 in the R programming language, optimized for measurements from plate readers (Boutros *et al*., 2006), and more recently cytomapper, an R package for analyzing imaging mass cytometry experiments (Eling *et al*., 2021). Of note, efforts are also ongoing to develop general-purpose HCS analysis tools in R and Python, with cytominer and Pycytominer, so far only available as GitHub repositories (Singh, 2021; Way, 2021). Despite the existence of these tools, the field still heavily relies on custom implementation of morphological profile curation for each study to account for different imaging modalities and analytical goals.

Julia is a high-performance, high-level open-source programming language specifically designed for scientific computing and data science (Bezanson *et al*., 2017). It is increasingly adopted by researchers in bioinformatics, with applications ranging from protein sequence analysis (Zea *et al*., 2016) to structural bioinformatics (Greener *et al*., 2020) and flux balance analysis (Heirendt *et al*., 2017). Julia is also ideal for tackling the challenges of morphological analyses, as they are both computationally demanding and inherently high-level. In this paper, we introduce BioProfiling.jl, the first Julia library for efficient and convenient morphological profiling that (i) handles noisy data through systematic filtering and robust statistics, (ii) provides dedicated functions to normalize data and mitigate layout effects and (iii) implements statistical tests for quantifying the strength of morphological changes that take the variability of morphological profiles into account. Our integrated software solution is thus bridging the existing gap between experimental data and biological interpretation. Furthermore, we conduct an image-based chemical screen to validate our approach and characterize the morphological impact of compounds in U-2-OS cells.

## Methods

### (1) Package implementation and features

We created BioProfiling.jl, a package for the Julia programming language that compiles over 30 methods and data structures for all steps in assembling and curating morphological profiles. To enable the bioimage analysis community to apply BioProfiling.jl to their own data, a complete documentation and a set of notebooks reproducing the analyses described in this paper are provided.

In brief, the whole process of morphological profiling is conceptually simplified by defining an *Experiment* object that includes both quantitative data and metadata in a tabular format, and methods able to interact with these objects directly to curate, transform and visualize the corresponding profiles. After creating the *Experiment* object from a table of morphological features, such as measurements obtained from CellProfiler (McQuin *et al*., 2018) or activation values from a deep neural network (Ando *et al*., 2017; Pawlowski *et al*., 2016; Lu *et al*., 2019), one would typically filter entries (rows representing biological units) and select features (columns representing phenotypic descriptors) with the *Filter* and *Selector* types respectively. Convenient shorthand is provided such as the *NameSelector* type to select features based on their name rather than their values, or the *CombinationFilter* type to join simple *Filter* objects with any logical operator. Profiles can then be transformed with the *logtransform!* and *normtransform!* methods, and *decorrelate!* discards highly correlated measurements. The filtered *Experiment* objects also support UMAP visualizations (Mcinnes *et al*., 2018) as implemented in UMAP.jl. Profiles can be visually inspected by highlighting images and individual cells matching a *Filter* with the *diagnostic_images* method. Finally, *robust_morphological_perturbation_value* and efficient implementations of statistical distances, described in detail below, are available for quantifying the significance of morphological changes induced by a particular perturbation.

Freely available from GitHub and the Julia package registry under the MIT license, BioProfiling.jl is part of a growing open-source software ecosystem ensuring that it stays flexible, maintainable and interoperable by encouraging contributions from external developers.

### (2) Cell culture

We selected the U-2-OS cell line as it is morphologically expressive and commonly used in HCS experiments (Rohban *et al*., 2017; Gustafsdottir *et al*., 2013; Wawer *et al*., 2014). U-2-OS cells (ATCC HTB-96) were cultured in high glucose DMEM (Thermo Fisher #11960044), 10% fetal bovine serum (Sigma Aldrich #F0804), 1x penicillin/streptomycin (Biowest #L0022-020) and 1mM sodium pyruvate (Thermo Fisher #11360070) and maintained in a humidified incubator (5% CO_2_, 37°C).

### (3) Chemical screen

A total of 311 compounds were selected to cover a wide range of biological processes and based on their propensity to impact cellular morphology in in-house and published studies (Wawer *et al*., 2014). A full list of the compounds and their concentration is provided in Supp. Table 1. Drugs were transferred to 384-well plates (PerkinElmer #6057302) using a liquid handler, in which 32 DMSO wells were used as a reference to assess the effect of the compounds as DMSO was used as solvent for the chemical library. Two drug plates were seeded in 50μL of culture medium with U-2-OS cells at 750 and 1500 cells per well, respectively, and incubated at 37 °C with 5% CO_2_ for 72h. Living cells were then washed three times with PBS and stained for 10 minutes using CellMask Orange Plasma membrane stain (Thermo Fisher #C10045). Cells were washed three more times with PBS and fixed with a solution of 4% Formaldehyde (Thermo Fisher #28908). After washing three more times with PBS, cells were permeabilized with 50 μL of permeabilization solution (PBS supplemented with 0.1x saponin-based permeabilization solution (Invitrogen #00-8333-56) and 5% FCS (Sigma #F7524)) for one hour. F-Actin was stained overnight with Phalloidin-488 staining solution (0.6 U/ml in permeabilization buffer; Thermo Fisher #A12379). Nucleic acids were stained with 30 μL of DAPI (5 ug/ml in PBS, Thermo Fisher #D1306) for 10 to 20 minutes. Finally, cells were washed three times with PBS, 50 μL of PBS solution was added per well and full wells were imaged (20 fields of view with a 20x LWD objective) on an Operetta High-Content Imaging System (PerkinElmer) using three fluorescence channels to detect DAPI (360-400 / 410-480 nm), Phalloidin (460-490 / 500-550 nm) and CellMask (520-550 / 560-630 nm).

### (4) Image analysis

We processed and analysed microscopy images using CellProfiler 3.1.8 (McQuin *et al*., 2018). In brief, the image quality was assessed, the intensities for experiments with high background noise were log-transformed, the illumination on each image was corrected based on background intensities before segmenting cell nuclei using global minimum cross entropy thresholding. Two successive secondary segmentation steps were performed using the propagation method (Jones *et al*., 2005) and global minimum cross entropy thresholding first on the CellMask then on the phalloidin channel to detect the cell bodies surrounding each nucleus. Finally, measurements were acquired for intensities in the nuclei and cytoplasms, granularity on all channels, textural and shape features, intensity distributions and number of neighboring cells less than 5 pixels away. This led to a total of 385 morphological features per cell.

### (5) Morphological profiling with BioProfiling.jl

All measurements were compiled for each cell in a BioProfiling.jl *Experiment* object, and non-numerical and uninformative features such as cell orientation were excluded from the profiles. We designed four cell filters to exclude technical outliers such as poorly segmented objects, which we set based on extreme feature values and validated visually with BioProfiling.jl diagnostic tools. These filters excluded cells with high CellMask to Phalloidin or DAPI to CellMask segmented area ratios, with a low nucleus form factor or with a high maximal CellMask intensity. We created median profiles for each field of view containing three valid cells or more. We then removed features which were constant across all DMSO controls or over the complete plate, log-transformed the values that were then centered and scaled per well on the median and median absolute deviation (MAD) of the control profiles in the matching row or column, in order to correct for plate layout effects. We then reduced redundancy in the profiles by ordering features by decreasing MAD and sequentially removing features with a Pearson’s correlation coefficient higher than 0.8 with any of the previously selected features. Lastly, we reduced the profiles to four dimensions with UMAP (Mcinnes *et al*., 2018), aiming to preserve the cosine distances between points, with *min_dist* set to 2 and all other parameters left to default values.

### (6) Hit detection with the robust Hellinger distance

BioProfiling.jl offers several statistical distances for quantifying the significance of morphological changes in HCS. The Mahalanobis distance takes the spread of the data in each dimension into account, which can be useful to compare two experimental conditions as previously described (Hutz *et al*., 2013). We also implemented the robust Mahalanobis distance which does not get biased by outliers by replacing the mean and covariance matrix by robust estimators of location and dispersion obtained using the minimum covariance determinant algorithm (Cabana *et al*., 2019; Rousseeuw and van Driessen, 1999). We already successfully used this approach in a HCS analysis (German *et al*., 2020). The profile filtering described in this analysis would be twice as compact using BioProfiling.jl and the nearly 100 lines dedicated to the quantification of morphological activity would be reduced to a one-liner and significantly accelerated thanks to parallelization and to the speed of Julia.

Note that the Mahalanobis distance is defined between a single point and a distribution. The Hellinger distance generalizes this concept for two distributions, by incorporating estimators of location and scatter of two distributions, and is defined as follows:

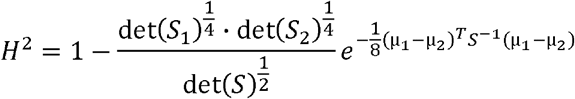

with *S* = (*S*_1_ + *S*_2_)/2, where *S*_1_, μ_1_ *S*_2_ and μ_2_ are the covariance matrices and means of the distributions 1 and 2, respectively. As for the robust Mahalanobis distance, we can substitute the covariance matrices *S*_1_ and *S*_2_ and the centers μ_1_ and μ_2_ using the Minimum Covariance Determinant (MCD) estimators and thus define the Robust Hellinger Distance (RHD) that we used to quantify the distance between DMSO controls and each chemical perturbation. One requirement for the MCD computation, and therefore for using the RHD, is to have twice as many measurements per condition as dimensions, which motivated our choice to work in a four-dimensional space in order to characterize most treatments. To assess the statistical significance of these values, we conducted a permutation test by shuffling the label of the points (perturbation or control) and calculating again the RHD 5000 times, which formed a null distribution associated with an empirical p-value. To accelerate this process, the permutations were computed in parallel by distributing computations on 16 threads. After Benjamini-Hochberg false discovery rate (FDR) correction, we obtained a significance score coined the Robust Morphological Perturbation Value (RMPV) and defined all compounds with an RMPV < 0.1, equivalent to an FDR cutoff of 10%, as morphological hits. Of note, the list of hits (Supp. Table 1.) was stable when doubling the number of permutations, showing that the process converged correctly.

### (7) Morphological and network distances

We integrated morphological profile information with publicly available data about each compound. First, we collected mechanisms of action (MOAs) and molecular targets from the LINCS perturbation database (Stathias *et al*., 2020). We queried the API for exact name matches or removed pharmaceutical salts or chirality when necessary to find the correct compound. All annotations are presented in Supp. Table 1. In total, 141 compounds had known targets and 112 were annotated with one or several MOAs. In particular, 23 MOAs were associated with 2 or more compounds and considered for downstream analysis. To compare morphological profiles between pairs of MOAs, we projected the profiles of the 59 hit compounds in four dimensions using UMAP and computed pairwise robust Hellinger distances as described above. The morphological distance between two MOAs was then defined as the average pairwise distance between compounds annotated to each MOA. We also obtained all human protein-protein interactions (PPIs) from the HIPPIE database (Alanis-Lobato *et al*., 2017), filtered out those with a confidence score below 0.63 (median of the score distribution), and assembled them into a PPI network. The conversion between gene symbols and ENTREZ identifiers of the targets was done with MyGene.info (Xin *et al*., 2016). We define the targets of a MOA as all known targets of the hit compounds associated with this MOA. We then assessed the network separation between the targets of each MOA using the s_AB_ score, which was previously found to be a good metric to study disease module and drug module separation (Menche *et al*., 2015; Caldera *et al*., 2019)]. The score is defined as:

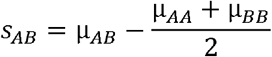

where μ_*AA*_ and μ_*BB*_ are the means of the minimum shortest network distances among the targets of MOA *A* and *B*, respectively, and μ_*AB*_ is the mean of the minimum shortest distance between the targets of MOA *A* and *B*.

### (8) Counting and classifying image artifacts

To quantify the prevalence of common imaging artifacts in HCS experiments, we visually inspected images from published studies deposited in the Image Data Resource (Breinig *et al*., 2015; Caldera *et al*., 2019; Rohban *et al*., 2017; Gustafsdottir *et al*., 2013; Williams *et al*., 2017). Depending on the study, we classified either all four fields of view from the wells A1 to A6 and B1 to B6 (Breinig *et al*., 2015; Caldera *et al*., 2019) or all nine fields of view from the wells A1 to A3 and B1 to B3 (Gustafsdottir *et al*., 2013; Rohban *et al*., 2017). Artifacts included dye clots and precipitations, cells not properly attached to the substrate, and other less frequent artifacts such as out-of-focus images or visible microwell edges. The most common artifacts only affected a restricted region of the image, suggesting that the unaffected parts of the images could be informative nonetheless, and motivating the extensive image filtering and quality controls performed in the respective studies. Given the overall abundance of such artifacts, however, we expect that a fraction of them will fail to be excluded and thus impact the image analysis and lead to outlier measurements.

## Results

### Profiling chemical perturbations with BioProfiling.jl

We conducted and analysed a chemical HCS experiment to study the morphological effect of small molecules in human osteosarcoma cells and demonstrate the applicability of BioProfiling.jl. In brief, we selected 311 compounds at a single concentration based on their morphological activity, and on their wide range of mechanisms of action (MOAs) and disease associations. U-2-OS cells were seeded on top of drug plates, and were fixed and stained to display nuclei, F-actin and total protein. Fluorescence images were acquired at a 20x magnification (Figure 2a). The images were analysed with CellProfiler (McQuin *et al*., 2018) and morphological descriptors were measured for each cell. These measurements were imported in Julia and used to define an *Experiment* object to be processed with BioProfiling.jl (Figure 2b). Two Jupyter notebooks enable the reproducibility of the following morphological profiling analysis (see Availability). First, filters are iteratively defined to identify cellular outliers based on extreme values. For instance, cells with unusually large cytoplasms compared to their nuclei were likely to be missegmented and therefore excluded (Figure 2c). After aggregating the profiles per image and discarding the least informative features, we reduced the dimensionality of the profiles to four dimensions using UMAP (Mcinnes *et al*., 2018). This formed a morphological space in which the profiles of some compounds, such as Vinblastine (tubulin inhibitor) and Wiskostatin (actin polymerization inhibitor) but also Pentamidine (antifungal agent), were clustered away from images of DMSO treatment (Figure 2d, Supp. Figure 1a). Using the dedicated methods for quantifying the significance of statistical distances implemented in BioProfiling.jl, we identified 248 compounds with a significant morphological activity compared to DMSO controls in a plate seeded with 750 cells per well (Figure 2e), coined morphological hits. In comparison, 242 hits were identified in a denser plate seeded with 1500 cells (Supp. Figure 1b). Of note, the seeding density had only a minor impact on whether compounds were identified as hits or not (Supp. Figure 1c). The hits on the two plates showed a large and highly significant overlap given the total number of tested compounds (Jaccard index of 0.78; *χ*^2^ test of independence: p = 1.5e-13).

**Fig. 2.**
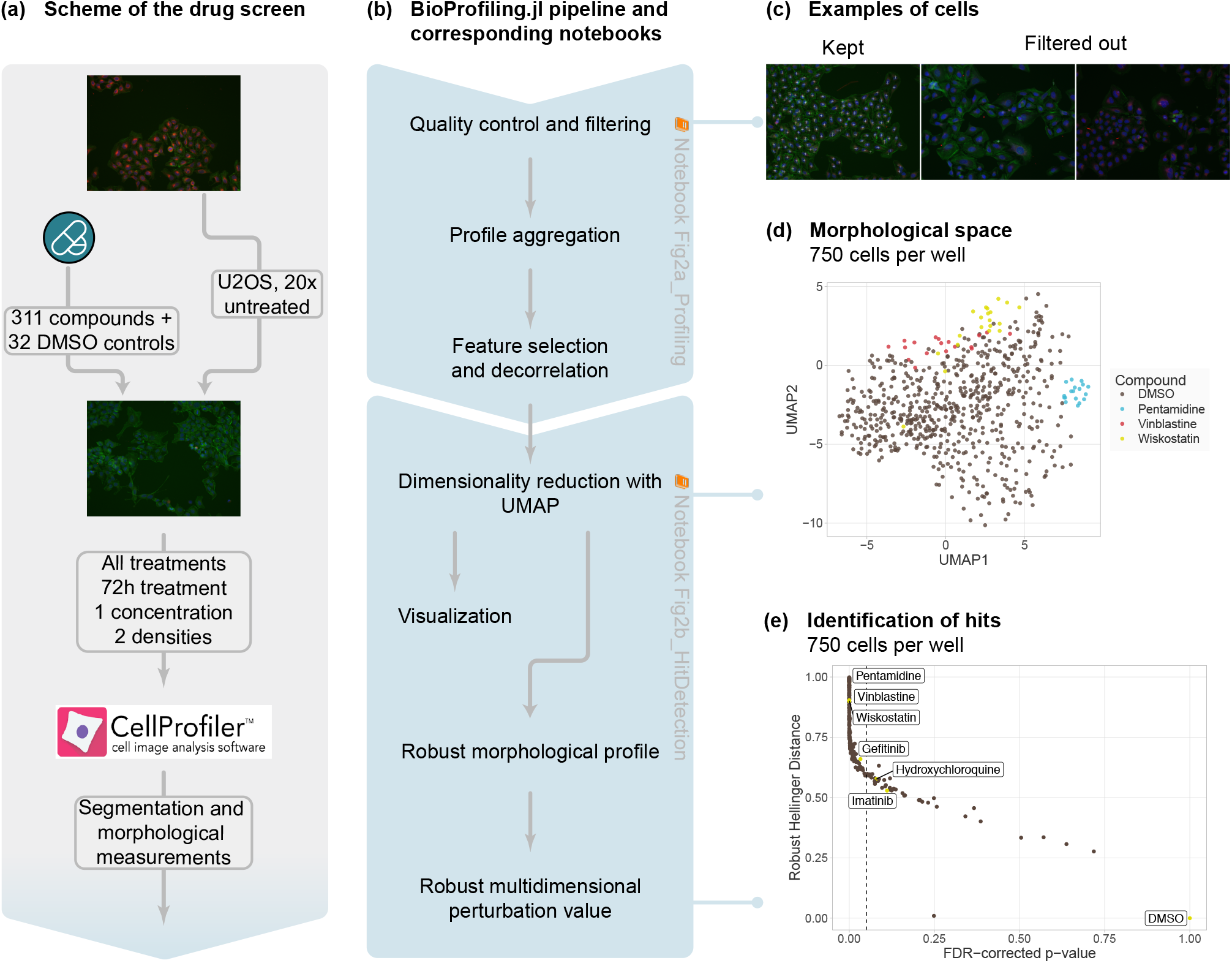
Robust cellular profiling with BioProfiling.jl characterizes the morphological diversity induced by pharmacologically-active compounds. (**a)** Experimental set-up of the HCS experiment. (**b)** Computational workflow using BioProfiling.jl. Boxes are annotated with the name of the notebooks with which to reproduce the analyses. (**c)** Example of images displaying cells kept in the analysis (left) or problematic cells discarded by one of the quality-control filters (center, right). Cytoplasm and nucleus centers are marked with a white cross for each cell. (**d)** UMAP embedding preserving the cosine distance between the morphological profiles aggregated per field of view in the plate seeded with 750 cells per well. Two out of four dimensions are represented. (**e)** Robust Hellinger distance and Robust Morphological Perturbation Value (FDR-corrected p-value) of each compound in the plate seeded with 750 cells per well compared to DMSO. Vertical dotted line indicates an FDR threshold of 0.1 and all compounds on its left are defined as morphological hits.

### Exploring mechanisms of action of active compounds

We next went on to characterize the compounds with a strong morphological impact identified on the plate seeded with 750 cells per well. Among the wide range of MOAs covered by the library, ten hit compounds were known Dopamine receptor antagonists, six hit compounds were annotated to Calcium channel blockers and six to Adrenergic receptor antagonists (Figure 3a). In this experiment, some MOAs were likelier than others to induce morphological changes, often in accordance to their biological role. In particular, all Tubulin inhibitors caused cytoskeletal defects and were identified as hits. In contrast, only half of the Topoisomerase inhibitors, which modulate DNA replication and transcription and are more likely to impact cell shape indirectly, if at all, were found to modulate the morphology. We note the presence of many oncological and chemotherapeutic agents (PDGFR receptor inhibitors, Topoisomerase inhibitors, KIT inhibitors, Tubulin inhibitors) and neurological drugs (Dopamine receptor antagonists, Tricyclic antidepressants, Norepinephrine reuptake inhibitors) among the morphological hits. Cell shape indeed plays an essential role in cancers, as cancerous cells are typically diagnosed by pathologists based on their morphology. Cell proliferation and several signaling pathways are also associated with cell geometry (Sero *et al*., 2015; Dupont *et al*., 2011; Aragona *et al*., 2013). Some compounds used to treat neurological disorders were also previously reported to induce morphological changes (Wawer *et al*., 2014), yet the mechanisms linking morphology and disease phenotype are still to be uncovered.

**Fig. 3.**
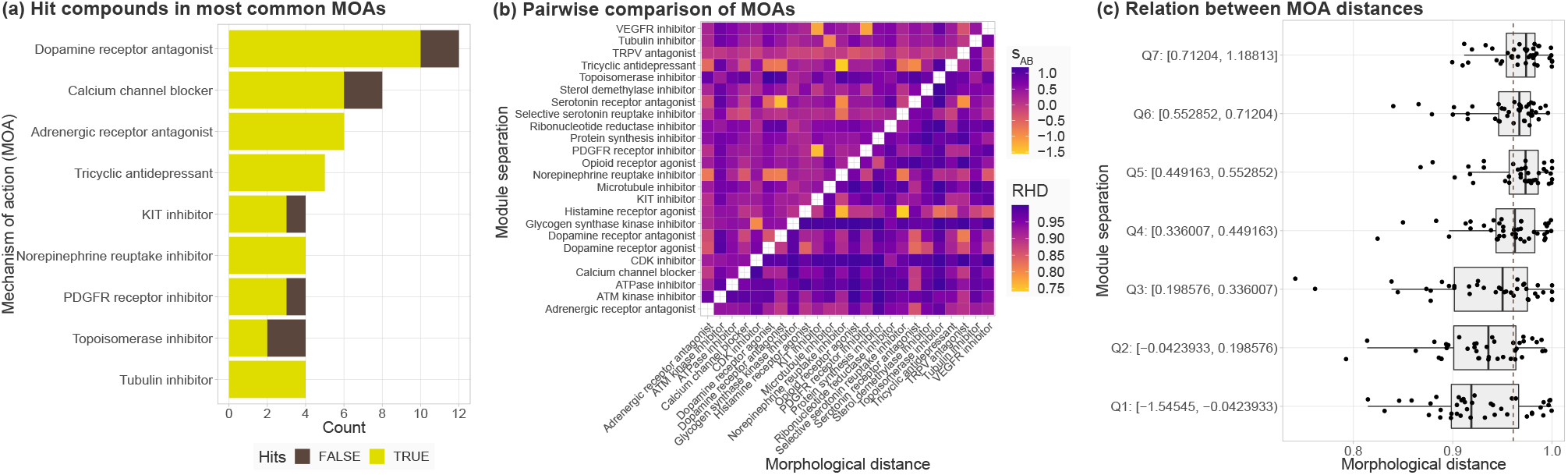
Morphological profiling and data integration characterize compound mechanisms of action (MOAs). **(a)** Number of hits and total number of compounds for the most common MOAs in the chemical library. **(b)** Dissimilarity of the molecular targets on a PPI network (s_AB_ score, upper triangle) and of the morphological profiles (RHD, lower triangle) for the MOAs with at least 2 hit compounds. **(c)** Relation between drug module separation (bins of s_AB_ scores) and morphological distance (RHD). RHD: Robust Hellinger Distance. PPI: Protein-Protein Interaction.

### Integrating target properties and morphological profiles

To further exploit the richness of the morphological profiles, we can go beyond quantifying whether they differ from untreated controls and also compare effects between compounds. We quantified the similarity between the morphological impact of MOAs by aggregating the mean of the pairwise robust Hellinger distance between their respective hit profiles (Figure 3b). While each MOA had a distinct signature, Glycogen synthase kinase inhibitors and CDK inhibitors were consistently distant from all other MOAs, hinting that modulation of kinase activity and cell signaling is likely to impact the cellular morphology in broad and distinctive ways as opposed to inducing a particular cytoskeletal defect.

While the morphological profiles are informative by themselves, they are best used by integrating additional information about the perturbations they describe. Here, we used available information on the targets of the compounds to contextualize their molecular environment within the protein-protein interaction network. We quantified the network separation between the targets of different MOAs via the s_AB_ score, which was used previously to quantify the separation of disease and drug modules (Menche *et al*., 2015; Caldera *et al*., 2019). A positive score is associated with well-separated sets of nodes, whereas a negative score corresponds to an overlap. We found that all the strongest network overlaps between MOAs corresponded to shared drug classes. Tricyclic antidepressants and Norepinephrine reuptake inhibitors corresponded to Non-selective monoamine reuptake inhibitors (ATC code N06AA). Serotonin receptor antagonists and Dopamine receptor antagonists both included Antipsychotics (ATC code N05A). PDGFR receptor inhibitors and KIT inhibitors were annotated to the exact same compound, Imatinib mesylate, Pazopanib and Sunitinib, which are all Protein kinase inhibitors (ATC code L01E).

When comparing morphological profiles, Histamine receptor antagonists were close to many other MOAs and showed the most striking similarities with Selective serotonin reuptake inhibitors and Norepinephrine reuptake inhibitors. Of note, Histamine receptor antagonists displayed a consistent level of network similarity with all other MOAs. The s_AB_ values close to zero reflect in part the spread of their 12 molecular targets on the PPI network, suggesting that generic PPI alteration patterns may correspond to morphological effects that are distinctive, but not unique.

By comparing morphological distances to molecular network separation, we observed that overlapping target modules are associated with more similar morphological profiles (Figure 3c). The effect does not fully explain morphological variability, which emphasizes the presence of intermediate regulatory processes between genotype and phenotype, and that the disruption of some biological processes are not detectable with general cell shape descriptors as experimental readouts. The quantification of morphological distances between profiles based on the UMAP-reduced space also means that part of the information contained in the original data is lost or distorted. Future development of methodologies leveraging the manifold learned by the UMAP method without the need for an embedding in Euclidean space will further alleviate this limitation. However, our results so far already confirm that there is a general association between the PPI network neighborhood targeted by a compound and their morphological outcome. This could be further explored to systematically link cellular morphology to function in health and disease.

## Discussion

HCS experiments offer a scalable and cost-efficient way to assess multiple conditions in a single experiment with a rich cellular readout. Assembling morphological profiles to describe these experimental conditions is thus essential and requires dedicated tools for data curation, feature selection, quality control, visualization and quantification of morphologically active perturbations. We implemented these tools in a single open-source software with intuitive and flexible data structures and syntax. We demonstrated by a concrete use case how BioProfiling.jl enables new research and allows the exploration of changes in cellular morphology by easing the analysis of large high-content imaging screens.

As Julia is an efficient programming language and allows parallelization of the computations, BioProfiling.jl can process large datasets in a performant manner. The biggest limitation for analysing large experiments at the single cell level is currently the memory usage, as the full set of morphological measurements needs to be loaded, which can be an issue on personal computers. This may be improved in the future by using lazy loading and allowing the user to process the data by batches. While we did not comment on individual feature changes in this study, the morphological activity of perturbations can be further characterized if the analysis relies on interpretable measurements. BioProfiling.jl can also help identify which features are varying the most and format the data for other tools, for instance to represent the typical cell morphology in a particular condition (Sailem *et al*., 2015; Khawatmi *et al*., 2021). Of note, by offering a systematic way to define filters for data curation and feature selection, BioProfiling.jl paves the way for data-driven feature engineering and machine-learning-powered artifact removal to further automate the process of morphological profiling.

BioProfiling.jl expands the existing landscape of resources available for biological data analysis, as illustrated in our application where we processed morphological measurements so that they can be integrated with protein-protein interactions as well as chemical annotations. The library contributes to the growing package ecosystem for bioinformatics in Julia which ensures that the morphological profiling analyses can be combined in larger projects together with other tasks ranging from sequencing to systems biology (Zea *et al*., 2016; Heirendt *et al*., 2017; Greener *et al*., 2020), and other libraries are conveniently available to integrate these different data types (Zakeri *et al*., 2018). Julia’s interoperability with other programming languages also makes the onboarding easy for users with prior programming experience who, for instance, might prefer to perform certain tasks in R or Python. This is demonstrated in the provided Jupyter notebooks, with all plots being generated using R’s ggplot2 library (Wickham, 2016) and with the computation of the MCD estimators for robust statistical distances which relies on R’s robustbase package, which in turn calls efficient Fortran routines.

Despite being initially designed and extensively tested for morphological profiling, the ability of BioProfiling.jl to handle large high-dimensional datasets and provide dedicated robust normalization and comparison methods could also be leveraged for other data analyses such as single cell transcriptomics or metabolomics experiments, which also require the curation and transformation of data in tabular format.

Finally, we conducted and analysed a chemical high-content imaging screen, exemplifying a concrete use case of the BioProfiling.jl package and characterizing the effect of small molecules across diverse MOAs. Since the compounds used in the chemical screen were initially selected to cover a wide range of morphological activity, the interpretation of MOA enrichment among hits is limited. While large-scale, hypothesis-free screening of small molecules can offer an unbiased view of the compound types that affect cellular morphology (Wawer *et al*., 2014; Bryce *et al*., 2019), our library design enabled us to observe significant changes induced by more than three quarters of the used compounds. We observed both commonalities and differences in the effect induced by different MOAs. The corresponding morphological profiles were further integrated with the information available about the PPI network properties of the compound targets, which proved to offer complementary views of compound effects and emphasized the role high-content screening could play in unraveling the relationship between cellular morphology and function.

## Acknowledgements

C.W.F is supported by a DOC-fellowship of the Austrian Academy of Sciences: 25525. We thank Raphael Bednarsky for discussion and feedback on the library and Daniel Malzl for his feedback on the manuscript.

## Funding

This work was supported by the Vienna Science and Technology Fund (WWTF) through project VRG15-005.

### Conflict of Interest

none declared.

## Notes

### Competing Interest Statement

The authors have declared no competing interest.

