## Supplementary Figure 1 and Supplementary Table 1 for "BioProfiling.jl: Profiling biological perturbations with high-content imaging in single cells and heterogeneous populations"

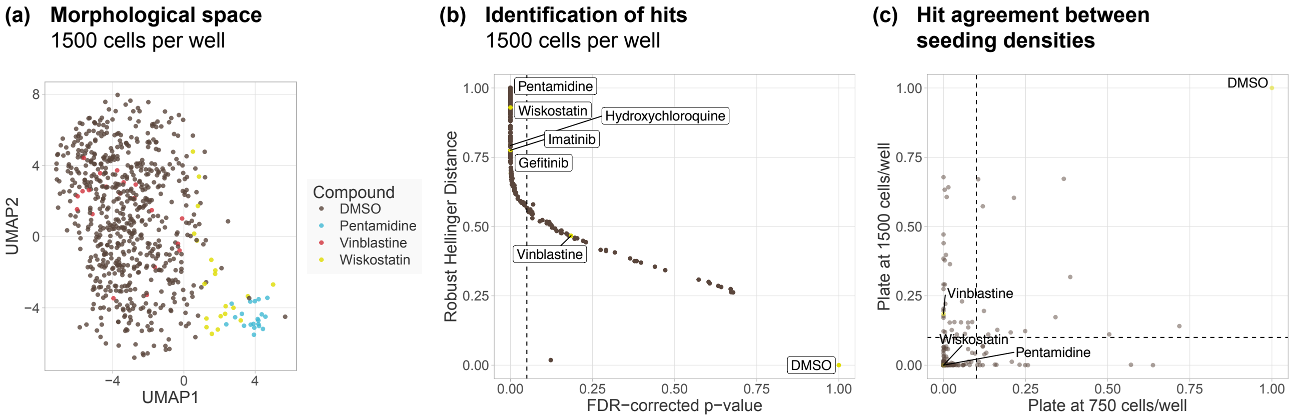


**Supp. Figure 1: BioProfiling.jl profiles of plates seeded at 750 and 1500 cells per well curated with are similar. (a)** UMAP embedding preserving the cosine distance between the mor-phological profiles aggregated per field of view in the plate seeded with 1500 cells per well. Two out of four dimensions are represented. **(b)** Robust Hellinger distance and Ro-bust Morphological Perturbation Value (FDR-corrected p-value) of each compound in the plate seeded with 1500 cells per well compared to DMSO. Vertical dotted line indicates an FDR threshold of 0.1 and all compounds on its left are defined as morphological hits. **(c)** FDR-corrected p-value of the significance of morphological changes induced by each compound in both plates. Dotted lines indicate an FDR threshold of 0.1.

| CompoundName | MOA | Targets | RMPV750 | RMPV1500 |
|---|---|---|---|---|
| (+)-Butaclamol hydrochloride | | | 0.2479179 | 0 |
| (+)-Cyclazocine | | | 0.0288018 | 0.0012478 |
| (+/-)-Sulfinpyrazone | ["Uricosuric blocker"] | ["ABCC1", "ABCC2", "FPR1", "SLC22A12"] | 0.0019172 | 0.015413 |
| (-)-JQ1 | | | 0.0003682 | 0 |
| (-)-Perillic acid | | | 0.1570008 | 0.0018222 |
| (-)-Quinpirole hydrochloride | ["Dopamine receptor agonist"] | ["DRD2", "DRD3", "DRD4", "DRD1", "HTR1A", "HTR2A", "HTR2B", "HTR2C"] | 0.0016163 | 0.1061314 |
| (-)-trans-(1S,2S)-U-50488 hydrochloride | | | 0.0087051 | 0.027791 |
| (S)-(+)-Camptothecin | | | | 0 |
| (S)-Propranolol hydrochloride | ["Adrenergic receptor antagonist"] | ["ADRB2", "ADRB3", "ADRB1", "CYP2C19", "HTR1A", "HTR1B"] | 0.0286805 | 0 |
| (±)-Isoproterenol hydrochloride | | | | |
| (±)-Methoxyverapamil hydrochloride | | | | |
| (±)-Metoprolol (+)-tartrate | | | | |
| (±)-Octoclothepin maleate | | | | |
| (±)-SKF-38393 hydrochloride | | | | |
| (±)-Sulpiride | | | | |
| (±)-Verapamil hydrochloride | | | | |
| (±)-alpha-Lipoic Acid | | | | |
| 1,10-Phenanthroline monohydrate | | | 0.020785 | 0.0026091 |
| 1,7-Dimethylxanthine | | | 0.0558376 | 0.1603626 |
| 2,2'-Bipyridyl | | | 0.0013317 | 0.0307685 |
| 2,3-Dimethoxy-1,4-naphthoquinone | | | 0.0055309 | 0.0642102 |
| 2-Phenylaminoadenosine | | | 0.1222335 | 0 |
| 2-methoxyestradiol | | | 0 | 0.1732954 |
| 4-(2-Aminoethyl)benzenesulfonyl fluoride hydrochloride | | | 0 | 0.0015266 |
| 4-Hydroxy-3-methoxyphenylacetic acid | | | 0 | 0.1914777 |
| 5-(N,N-hexamethylene)amiloride | | | 0 | 0 |
| 5-(N-Ethyl-N-isopropyl)amiloride | | | 0 | 0 |
| 5-Bromo-2'-deoxyuridine | | | 0 | 0 |
| 5-Fluorouracil | | | 0 | 0 |
| 5-azacytidine | | | 0 | 0 |
| 5HPP-33 | | | 0.3979575 | 0.3161221 |
| 5alpha-Pregnan-3alpha-ol-20-one | | | 0.0173094 | 0 |
| 6,7-ADTN hydrobromide | | | 0 | 0.3030974 |
| 6-Nitroso-1,2-benzopyrone | | | 0 | 0 |

| CompoundName | MOA | Targets | RMPV750 | RMPV1500 |
|---|---|---|---|---|
| 7-Cyclopentyl-5-(4-phenoxy)phenyl-7H-pyrrolo[2,3-d]pyrimidin-4-ylamine | | | 0.0016163 | 0 |
| A-77636 hydrochloride | | | | |
| AC-93253 iodide | | | 0 | 0 |
| AEG 3482 | | | 0 | 0 |
| AMG 9810 | ["TRPV antagonist"] | ["TRPV1"] | 0.0314152 | 0.000328 |
| AZ191 | | | | |
| Acepromazine maleate | ["Dopamine receptor antagonist"] | ["ADRA1A", "ADRA1B", "DRD1", "DRD2", "HTR1A", "HTR2A"] | 0.1114224 | 0.0096587 |
| Adaphostin | | | | |
| Albendazole | ["Anthelmintic", "Tubulin inhibitor"] | ["CYP1A2", "CYP2J2", "TUBA1A", "TUBB", "TUBB4B"] | 0 | 0 |
| Aliskiren | ["Antihypertensive", "Peptidase inhibitor", "Protease inhibitor", "Renin inhibitor"] | ["REN"] | 0.0063378 | 0.2233888 |
| Ammonium pyrrolidinedithiocarbamate | | | | 0 |
| Amodiaquine | ["Histamine receptor agonist"] | ["HNMT", "CYP2C8"] | 0 | 0 |
| Amsacrine hydrochloride | ["Topoisomerase inhibitor"] | ["TOP2A", "KCNH2"] | 0.2100677 | 0.58589 |
| Ancitabine hydrochloride | | | 0 | 0 |
| Apomorphine hydrochloride hemihydrate | | ["DRD2", "DRD5", "ADRA2A", "ADRA2B", "ADRA2C", "DRD1", "DRD3", "DRD4", "HTR1A", "HTR2A", "HTR2B", "HTR2C", "TRPA1", "CALY", "HTR1B", "HTR1D"] | 0.0125635 | 0.0012478 |
| Arbidol hydrochloride | | | 0.0039283 | 0.1244432 |
| Auranofin | | | | |
| Aurora-A Inhibitor I | | | 0.122102 | 0.0731426 |
| Aurothioglucose | | | 0 | |
| Azatadine | | | 0.1125291 | 0.6572704 |
| Azithromycin | ["Bacterial 50S ribosomal subunit inhibitor"] | ["MLNR"] | 0.1358873 | 0.1024149 |
| BAY 61-3606 hydrochloride hydrate | | | | 0 |
| BIO | | | 0 | 0.000328 |
| BIX 01294 trihydrochloride hydrate | | | | |
| BMS-193885 | | | | 0 |
| BRD3308 | | | 0 | 0 |
| BTO-1 | | | 0.0841943 | 0.0117533 |
| BW 723C86 | ["Serotonin receptor agonist"] | ["HTR2B", "HTR2A", "HTR2C"] | 0.0125635 | 0 |
| Bay 11-7082 | | | | |
| Bay 11-7085 | | | | |
| Benidipine hydrochloride | ["Calcium channel blocker"] | ["CACNA1C", "CACNA1G", "CYP3A5"] | 0 | 0 |
| Benoxathian hydrochloride | | | 0.0003682 | 0 |

| CompoundName | MOA | Targets | RMPV750 | RMPV1500 |
|---|---|---|---|---|
| Benzamil hydrochloride | | ["PKD2L1", "SCNN1A", "SCNN1B", "SCNN1G", "ASIC1", "SCNN1D", "SLC8A1"] | 0.0003682 | 0 |
| Benztropine mesylate | | | 0 | 0 |
| Bortezomib | | | | |
| Brefeldin A from Penicillium brefeldianum | ["Protein synthesis inhibitor", "Brefeldin A inhibited guanine nucleotide exchange protein inhibitor", "Golgi-specific brefeldin A-resistance guanine nucleotide exchange factor inhibitor"] | ["ARF1", "ARFGEF1", "ARFGEF2", "CYTH2", "GBF1", "SAR1A"] | 0 | 0 |
| Brequinar sodium salt hydrate | | | 0 | 0 |
| Budesonide | | ["NR3C1", "CYP3A5", "CYP3A7"] | 0 | 0 |
| CCCI-01 | | | 0 | 0 |
| CCT137690 | | | | 0 |
| CGS-15943 | ["Adenosine receptor antagonist"] | ["ADORA1", "ADORA2A", "ADORA2B", "ADORA3"] | 0.1351209 | 0.5604215 |
| CID 11210285 hydrochloride | | | 0 | 0 |
| CID2858522 | | | 0.0052674 | 0 |
| CP466722 | ["ATM kinase inhibitor"] | ["ATM"] | 0 | 0 |
| CYM50358 | | | 0 | 0 |
| Caffeic Acid | ["Lipoxygenase inhibitor", "HIV integrase inhibitor", "NFkB pathway inhibitor", "Nitric oxide production inhibitor", "PPAR receptor modulator", "TNF production inhibitor", "Tumor necrosis factor production inhibitor"] | ["ALOX5", "MIF", "RELA", "TNF"] | 0 | 0.0062207 |
| Caffeic acid phenethyl ester | | ["RELA"] | 0.0326187 | 0 |
| Calcimycin | | | | |
| Cantharidic Acid | | | 0.0070918 | 0.0096587 |
| Cantharidin | | String[] | 0.0052674 | 0 |
| Carmofur | ["Thymidylate synthase inhibitor"] | ["TYMS"] | 0 | 0 |
| Carvedilol | ["Adrenergic receptor antagonist"] | ["ADRB1", "ADRB2", "ADRA1A", "ADRA1B", "ADRA1D", "ADRA2A", "ADRA2B", "ADRA2C", "ADRB3", "CYP2C19", "CYP2E1", "GJA1", "HIF1A", "KCNH2", "NDUFC2", "NPPB", "RYR2", "SELE", "VCAM1", "VEGFA"] | 0 | 0 |
| Cerivastatin | | | | |
| Chloroquine | ["Antimalarial"] | ["CYP2C8", "GSTA2", "MRGPRX1", "TLR9", "TNF"] | 0 | 0 |

| CompoundName | MOA | Targets | RMPV750 | RMPV1500 |
|---|---|---|---|---|
| Chlorpromazine hydrochloride | ["Dopamine receptor antagonist"] | ["DRD2", "ADRA2A", "ADRA2B", "ADRA2C", "DRD1", "DRD3", "DRD4", "DRD5", "HRH1", "HTR1A", "HTR2A", "HTR2C", "HTR6", "HTR7", "ADRA1A", "ADRA1B", "ADRA1D", "CALM1", "CHRM1", "CHRM3", "HRH4", "HTR2B", "KCNH2", "KIF11", "ORM1", "ORM2", "SMPD1", "TRPC5"] | 0.0022114 | 0.6397159 |
| Chlorprothixene hydrochloride | ["Dopamine receptor antagonist"] | ["DRD2", "CHRM1", "CHRM2", "CHRM3", "CHRM4", "CHRM5", "DRD1", "DRD3", "HRH1", "HTR2A", "HTR2B", "HTR2C"] | 0 | 0 |
| Cilengitide trifluoroacetic acid salt | | | 0 | 0 |
| Cilnidipine | ["Calcium channel blocker"] | ["CACNA1B", "CACNA1C"] | 0.0007038 | 0 |
| Cinacalcet | | | | |
| Clemizole hydrochloride | | String[] | 0.2635852 | 0 |
| Clodronic acid | | ["SLC25A4", "SLC25A5", "SLC25A6"] | 0.0010296 | 0.0635064 |
| Clofarabine | ["Ribonucleoside reductase inhibitor"] | ["RRM1", "POLA1", "RRM2", "SLC22A8"] | 0.0332124 | 0 |
| Clomipramine hydrochloride | ["Serotonin transporter inhibitor (SERT)"] | ["SLC6A4", "SLC6A2", "CYP2C19", "GSTP1", "HTR2A", "HTR2B", "HTR2C", "SLC6A3"] | 0.1222335 | 0.0409639 |
| Clotrimazole | ["Cytochrome P450 inhibitor", "Imidazoline receptor ligand"] | ["KCNN4", "CYP3A4", "CYP51A1", "NR1I2", "NR1I3", "TRPM2", "TRPM4", "TRPM8"] | 0 | 0 |
| Colchicine | | ["TUBB", "GLRA1", "GLRA2", "TUBB1"] | 0.01668 | 0.6354709 |
| Cyclosporin A | | ["PPIA", "ABCB11", "CAMLG", "CYP3A5", "CYP3A7", "FPR1", "PPID", "PPIF", "PPP3CA", "PPP3R2", "SLC10A1", "SLCO1B1", "SLCO1B3"] | 0 | 0 |
| Cyproterone acetate | ["Androgen receptor antagonist", "Progesterone receptor agonist", "Testosterone receptor antagonist"] | ["AR", "ADORA1", "ESR1"] | 0 | 0.0178228 |
| Cytarabine | ["Ribonucleotide reductase inhibitor"] | ["POLB", "POLA1"] | 0 | 0 |
| Cytosine-1-beta-D-arabinofuranoside hydrochloride | | | | 0 |
| D-Cycloserine | ["Bacterial cell wall synthesis inhibitor"] | ["GRIN1"] | 0.0790705 | 0.0635064 |
| D-ribofuranosylbenzimidazole | | | 0.0044839 | 0.000328 |
| DCEBIO | ["Potassium channel activator"] | ["KCNN2", "KCNN3", "KCNN4"] | 0 | 0.686793 |
| Danshensu sodium salt | | | 0.0039283 | 0.2793182 |
| Dantrolene sodium | ["Calcium channel blocker"] | ["RYR1", "RYR3"] | 0.6349681 | 0 |
| Dequalinium chloride hydrate | | ["KCNN1", "KCNN3"] | 0 | 0 |

| CompoundName | MOA | Targets | RMPV750 | RMPV1500 |
|---|---|---|---|---|
| Digitoxin | | | | |
| Dihydroartemisinin | | | 0 | 0 |
| Dihydroergotamine | | ["HTR1D", "HTR1B", "ADRA2A", "DRD2", "HTR1E", "HTR1F", "HTR2B", "HTR7"] | 0.1428492 | 0.0400524 |
| Dihydroergotamine methanesulfonate | | ["HTR1D", "HTR1B", "ADRA2A", "DRD2", "HTR1E", "HTR1F", "HTR2B", "HTR7"] | 0.0034168 | 0.0070343 |
| Dihydroouabain | | | 0.0052674 | 0 |
| Diphenyleneiodonium chloride | ["Nitric oxide synthase inhibitor"] | ["ALDH1A2", "ALDH2", "ALDH5A1", "ALDH7A1", "NOX3", "XDH"] | 0 | 0 |
| Dipyridamole | ["Phosphodiesterase inhibitor"] | ["ADA", "PDE5A", "PDE10A", "PDE4A", "PDE7B", "PDE8A", "PDE8B", "SLC29A1"] | 0.3587426 | 0.6785284 |
| Docetaxel | | ["TUBB", "BCL2", "MAP2", "MAP4", "MAPT", "NR1I2", "TUBB1"] | 0 | 0 |
| Domperidone | ["Dopamine receptor antagonist"] | ["DRD2", "DRD3", "ABCG2", "CYP3A5"] | 0 | 0 |
| Doxazosin mesylate | ["Adrenergic receptor antagonist"] | ["ADRA1D", "ADRA1A", "ADRA1B", "CYP2C19", "KCNH2", "KCNH6", "KCNH7"] | 0.0106818 | 0.6036192 |
| Doxorubicin | | ["TOP2A"] | 0 | 0 |
| Doxycycline hydrochloride | ["Bacterial 30S ribosomal subunit inhibitor", "Metalloproteinase inhibitor"] | ["MMP8", "MMP1"] | 0.0003682 | 0 |
| Droperidol | ["Dopamine receptor antagonist"] | ["DRD2", "ADRA1A"] | 0.0044839 | 0.394313 |
| E-64 | | | 0 | 0.1732954 |
| Ebastine | | | 0 | 0 |
| Efavirenz | ["HIV protease inhibitor"] | ["CYP2B6", "CYP2C19", "CYP2C8", "CYP3A4", "CYP3A5"] | 0.1994439 | 0.0140131 |
| Ellipticine | ["Topoisomerase inhibitor"] | ["TOP2A", "TOP2B"] | 0 | 0 |
| Emetine dihydrochloride hydrate | ["Protein synthesis inhibitor"] | ["RPS2"] | 0 | 0 |
| Enclomiphene hydrochloride | | | | |
| Endoxifen | | | | 0 |
| Eptifibatide | | | 0 | 0.1203325 |
| Ethinyl Estradiol | ["DNA directed DNA polymerase stimulant", "Estrogenic component in oral contraceptives", "Estrogen receptor agonist"] | ["CYP2C8", "ESR1", "ESR2", "NR1I2"] | 0.0957928 | 0.0478751 |
| Etoposide | | ["TOP2A", "CYP2E1", "CYP3A5", "TOP2B"] | 0 | 0 |
| Flunarizine dihydrochloride | ["Calcium channel blocker"] | ["CACNA1G", "CACNA1H", "CACNA1I", "CALM1", "CYP2J2", "HRH1"] | 0.0111745 | 0 |
| Fluoxetine hydrochloride | ["Selective serotonin reuptake inhibitor (SSRI)"] | ["SLC6A4", "ANO1", "CYP2C19", "HTR2B"] | 0 | 0 |

| CompoundName | MOA | Targets | RMPV750 | RMPV1500 |
|---|---|---|---|---|
| Fluspirilene | ["Dopamine receptor antagonist"] | ["DRD2", "HTR2A", "CACNG1", "HRH1", "HTR1A", "HTR1D", "HTR1E"] | 0.0206216 | 0 |
| Forskolin | | ["ADCY2", "ADCY5", "GNAS"] | 0.0013317 | 0.000328 |
| Furamidine dihydrochloride | | | 0 | 0 |
| GANT61 | | | 0 | 0.000328 |
| GBR-12909 dihydrochloride | | | 0 | 0 |
| GSK-650394 | | ["SGK1", "SGK2"] | 0 | 0.010711 |
| GSK1210151A | | | 0 | 0 |
| GW2974 | | | 0 | 0.0094738 |
| GW9662 | | | 0.0047572 | 0 |
| Gefitinib | ["EGFR inhibitor"] | ["EGFR", "CYP2C19"] | 0.0403039 | 0.0023549 |
| Gemcitabine hydrochloride | ["Ribonucleotide reductase inhibitor"] | ["RRM1", "CMPK1", "RRM2", "TYMS"] | 0 | 0 |
| Histamine, R(-)-alpha-methyl-, dihydrochloride | | | 0.5131839 | 0.1117667 |
| Hydroquinone | | | 0.0728151 | 0.1129409 |
| Hydroxychloroquine | | ["TLR7", "TLR9"] | 0.0762777 | 0 |
| IKK-16 dihydrochloride | ["IKK inhibitor"] | ["IKBKB"] | 0 | 0 |
| IMS2186 | | | 0 | 0 |
| IN-1130 | | | 0 | 0 |
| Icaritin | | | 0 | 0.4266664 |
| Idarubicin | | ["TOP2A"] | 0 | 0 |
| Idazoxan hydrochloride | | ["NISCH"] | 0 | 0 |
| Imatinib | ["BCR-ABL kinase inhibitor", "KIT inhibitor", "PDGFR receptor inhibitor"] | ["ABL1", "KIT", "PDGFRA", "BCR", "CSF1R", "PDGFRB", "ABCG2", "CYP2C19", "CYP2C8", "CYP3A5", "DDR1", "NTRK1", "RET"] | 0.1198048 | 0 |
| Imatinib mesylate | ["BCR-ABL kinase inhibitor", "KIT inhibitor", "PDGFR receptor inhibitor"] | ["ABL1", "KIT", "PDGFRA", "BCR", "CSF1R", "PDGFRB", "ABCG2", "CYP2C19", "CYP2C8", "CYP3A5", "DDR1", "NTRK1", "RET"] | 0.0039283 | 0 |
| Imipramine hydrochloride | ["Norepinephrine reuptake inhibitor", "Serotonin reuptake inhibitor"] | ["SLC6A2", "SLC6A4", "CHRM2", "ADRA1A", "ADRA1B", "ADRA1D", "CHRM1", "CHRM3", "CHRM4", "CHRM5", "CYP2C19", "DRD1", "DRD2", "DRD5", "HRH1", "HTR1A", "HTR2A", "HTR2C", "HTR6", "HTR7", "KCND2", "KCND3", "KCNH1", "KCNH2", "SLC6A3"] | 0 | 0 |
| Iodoacetamide | | | 0 | 0 |
| Irinotecan | | | | 0 |
| Isoproterenol | | | 0.122102 | 0.0733176 |
| JFD00244 | | | 0 | 0 |
| JS-K | | String[] | 0.0065918 | 0.0012478 |
| K114 | | | 0 | 0 |
| KB-R7493 | | | 0 | 0 |

| CompoundName | MOA | Targets | RMPV750 | RMPV1500 |
|---|---|---|---|---|
| KT203 | | | 0 | 0.0124547 |
| KU-55933 | ["ATM kinase inhibitor"] | ["ATM", "PRKDC"] | 0.0016163 | 0.0009729 |
| KY-05009 | | | 0 | 0 |
| Kenpaullone | ["CDK inhibitor", "Glycogen synthase kinase inhibitor"] | ["GSK3B", "CDK1", "CDK5", "CCNB1", "CDK2", "LCK"] | 0 | 0 |
| Ketoconazole | ["Sterol demethylase inhibitor"] | ["AR", "CYP19A1", "CYP21A2", "CYP2C19", "CYP3A5", "CYP3A7", "KCNA10"] | 0.0013317 | 0 |
| Ketotifen fumarate | ["Histamine receptor agonist", "Histamine receptor ligand", "Leukotriene receptor antagonist", "Phosphodiesterase inhibitor"] | ["HRH1", "PDE4A", "PDE4B", "PDE4C", "PDE4D", "PDE7A", "PDE7B", "PDE8A", "PDE8B", "PGD"] | 0.0063378 | 0.3894851 |
| L-703,606 oxalate salt hydrate | | | 0.0861122 | 0.1679892 |
| L-741,626 | | | 0 | 0 |
| L-Cycloserine | | | 0.072975 | 0.1591953 |
| L-Tryptophan | | | 0.0007038 | 0.0353625 |
| LDN-214117 | | | 0.0260147 | 0.0084 |
| LP 12 hydrochloride hydrate | | | 0.0007038 | 0 |
| LP44 | | | 0 | 0 |
| LY-294,002 hydrochloride | | | 0 | 0 |
| Lasofoxifene tartrate | | | | |
| Lercanidipine hydrochloride hemihydrate | | | 0.0019172 | 0 |
| Levetiracetam | ["Calcium channel blocker"] | ["SV2A", "CACNA1B", "SCN1A"] | 0.0264037 | 0.1362111 |
| Loperamide | ["Opioid receptor agonist"] | ["OPRM1", "OPRD1", "CACNA1A", "CALM1", "CYP2B6", "CYP2C8", "NPR2", "OPRK1", "POMC"] | 0 | 0 |
| Loperamide hydrochloride | ["Opioid receptor agonist"] | ["OPRM1", "OPRD1", "CACNA1A", "CALM1", "CYP2B6", "CYP2C8", "NPR2", "OPRK1", "POMC"] | 0 | 0 |
| Loratadine | ["Histamine receptor antagonist"] | ["HRH1", "CYP2C19", "CYP3A5"] | 0.1217486 | 0.1974474 |
| Lorcainide hydrochloride | | | 0 | 0.0020819 |
| Lubeluzole dihydrochloride | | | 0 | 0 |
| M-110 | | | 0 | 0 |
| MG 624 | | | 0 | 0 |
| MK-677 | | | 0 | 0 |
| ML-7 | ["Myosin light chain kinase inhibitor"] | ["MYLK"] | 0.0016163 | 0 |
| ML240 | | | 0.0882397 | 0.0403832 |
| ML324 | | | 0 | 0 |
| Maprotiline | ["Norepinephrine reuptake inhibitor", "Tricyclic antidepressant"] | ["SLC6A2", "ADRA1A", "ADRA1B", "ADRA1D", "ADRA2A", "ADRA2B", "ADRA2C", "CHRM1", "CHRM2", "CHRM3", "CHRM4", "CHRM5", "DRD2", "HRH1", "HTR2A", "HTR2C", "HTR7"] | 0.0395053 | 0.0015266 |

| CompoundName | MOA | Targets | RMPV750 | RMPV1500 |
|---|---|---|---|---|
| Maprotiline hydrochloride | ["Norepinephrine reuptake inhibitor", "Tricyclic antidepressant"] | ["SLC6A2", "ADRA1A", "ADRA1B", "ADRA1D", "ADRA2A", "ADRA2B", "ADRA2C", "CHRM1", "CHRM2", "CHRM3", "CHRM4", "CHRM5", "DRD2", "HRH1", "HTR2A", "HTR2C", "HTR7"] | 0 | 0 |
| Metergoline | | | | |
| Methiothepin mesylate | | | 0 | 0 |
| Methoxamine hydrochloride | | ["ADRA1A", "ADRA1B", "ADRA1D"] | 0.0013317 | 0.0536727 |
| Metrazoline oxalate | | | 0 | 0 |
| Mibefradil dihydrochloride | ["T-type calcium channel blocker"] | ["CACNA1G", "CACNA1H", "CACNA1C", "CACNA1I", "ANO1", "CACNA1D", "CACNA1F", "CACNA1S", "CACNB1", "CACNB2", "CACNB3", "CACNB4", "CATSPER1", "CATSPER2", "CATSPER3", "CATSPER4", "CYP3A5", "CYP3A7", "SCN2A", "SCN4A", "SCN5A", "SCN9A"] | 0 | 0 |
| Mifepristone | ["Glucocorticoid receptor antagonist", "Progesterone receptor antagonist"] | ["PGR", "NR3C1", "AR", "CYP2B6", "CYP2C8", "CYP3A5", "CYP3A7", "NR1I2"] | 0.0726538 | 0.004879 |
| Mitotane | ["Antineoplastic"] | ["CYP11B1", "CYP11A1", "CYP3A4", "ESR1", "FDX1"] | 0.0164938 | 0.1653031 |
| Mitoxantrone | ["Topoisomerase inhibitor"] | ["TOP2A", "PIM1"] | 0.7197892 | 0.151491 |
| Mycophenolic Acid | ["Dehydrogenase inhibitor", "Inositol monophosphatase inhibitor"] | ["IMPDH1", "IMPDH2"] | 0 | 0 |
| N-p-Tosyl-L-phenylalanine chloromethyl ketone | | | 0.0003682 | 0.0015266 |
| NG-Monomethyl-L-arginine acetate | | | 0.0577294 | 0 |
| Nestorone | | | 0.0007038 | 0.0593708 |
| Nicardipine hydrochloride | ["Calcium channel blocker"] | ["CACNA1C", "ADORA3", "ADRA1A", "ADRA1B", "ADRA1D", "CACNA1D", "CACNA2D1", "CACNB2", "CALM1", "CHRM1", "CHRM2", "CHRM3", "CHRM4", "CHRM5", "PDE1A", "PDE1B"] | 0 | 0.0763732 |
| Niclosamide | ["DNA replication inhibitor", "STAT inhibitor"] | ["STAT3"] | 0.0457316 | 0 |
| Nisoldipine | ["Calcium channel blocker"] | ["CACNA1C", "CACNA1D", "CACNA1S", "CACNA2D1", "CACNB2", "CYP3A5"] | 0 | 0 |
| Nitidine chloride | | | | |
| Nocodazole | ["Tubulin inhibitor"] | ["HPGDS"] | 0 | 0 |

| CompoundName | MOA | Targets | RMPV750 | RMPV1500 |
|---|---|---|---|---|
| Nortriptyline hydrochloride | ["Tricyclic antidepressant"] | ["KCNJ10", "SLC6A2", "SLC6A4", "ADRA1A", "ADRA1B", "ADRA1D", "ADRA2A", "ADRA2B", "ADRA2C", "ADRB1", "ADRB2", "ADRB3", "CHRM1", "CHRM2", "CHRM3", "CHRM4", "CHRM5", "CYP2C19", "DRD2", "HRH1", "HTR1A", "HTR2A", "HTR2C", "HTR6", "PGRMC1", "PIK3CD", "SIGMAR1"] | 0 | 0 |
| Olanzapine | ["Dopamine receptor antagonist", "Serotonin receptor antagonist"] | ["DRD2", "HTR2A", "HTR2C", "DRD1", "DRD3", "DRD4", "HRH1", "HTR1A", "HTR1B", "HTR1D", "HTR1E", "HTR6", "HTR7", "ADRA1A", "ADRA1B", "ADRA2A", "ADRA2B", "ADRA2C", "ADRB1", "ADRB2", "ADRB3", "CHRM1", "CHRM2", "CHRM3", "CHRM4", "CHRM5", "CYP2C8", "DRD5", "GABRA1", "GABRA2", "GABRA3", "GABRA4", "GABRA5", "GABRA6", "GABRB1", "GABRB2", "GABRB3", "GABRD", "GABRE", "GABRG1", "GABRG2", "GABRG3", "GABRP", "GABRQ", "HRH2", "HRH4", "HTR1F", "HTR2B", "HTR3A", "HTR5A"] | 0.0076203 | 0.0777292 |
| Ouabain | ["ATPase inhibitor"] | ["ATP1A1", "ATP1A2", "ATP1A3", "ATP1A4", "ATP1B1", "ATP1B2", "ATP1B3", "ATP1B4", "FXYD2"] | 0 | 0 |
| PAPP | | | 0 | 0 |
| PD-407824 | | | | 0 |
| PD153035 hydrochloride | | | 0.0068431 | 0.0056832 |
| PF-429242 dihydrochloride | | | 0 | 0 |
| PMEG hydrate | | | | 0 |
| Palonosetron hydrochloride | ["Serotonin receptor antagonist"] | ["HTR3A"] | 0 | 0.0040382 |
| Paroxetine hydrochloride hemihydrate (MW = 374.83) | ["Selective serotonin reuptake inhibitor (SSRI)"] | ["SLC6A4", "CHRM1", "CHRM2", "CHRM3", "CHRM4", "CHRM5", "HTR2A", "SLC6A2"] | 0 | 0 |
| Parthenolide | | | | |
| Pazopanib | ["KIT inhibitor", "PDGFR receptor inhibitor", "VEGFR inhibitor"] | ["KDR", "KIT", "FLT1", "FLT4", "PDGFRB", "PDGFRA", "BRAF", "CSF1R", "CYP2B6", "CYP2C8", "CYP2E1", "DDR2", "FGF1", "FGFR1", "FGFR3", "ITK", "SH2B3"] | 0.0037067 | 0 |
| Pentamidine | | ["TRDMT1"] | 0 | 0 |
| Pentamidine isethionate | | ["TRDMT1"] | 0 | |

| CompoundName | MOA | Targets | RMPV750 | RMPV1500 |
|---|---|---|---|---|
| Pergolide methanesulfonate | ["Dopamine receptor agonist"] | ["DRD1", "DRD2", "ADRA2A", "ADRA2B", "ADRA2C", "DRD3", "DRD4", "DRD5", "HTR1A", "HTR1B", "HTR1D", "HTR2A", "HTR2B", "HTR2C", "ADRA1A", "ADRA1B", "ADRA1D", "KCNA5"] | 0 | 0.0511609 |
| Perphenazine | ["Dopamine receptor antagonist"] | ["DRD2", "CALM1", "DRD1", "HRH1", "HTR2A", "HTR2C", "HTR6", "HTR7"] | 0.0039283 | 0.0124547 |
| Phenamil methanesulfonate | ["TRPV antagonist"] | ["PKD2L1"] | 0.0325161 | 0.1496016 |
| Pheniramine maleate | ["Histamine receptor antagonist"] | ["HRH1"] | 0.0168645 | 0.0023549 |
| Phorbol 12-myristate 13-acetate | ["PKC activator"] | ["CD4", "KCNT2", "PRKCA", "TRPV4"] | 0 | 0 |
| Pifithrin-mu | ["HSP inhibitor"] | ["HSPA1A", "TP53"] | 0 | 0.000328 |
| Pimozide | ["Dopamine receptor antagonist"] | ["DRD2", "DRD3", "CACNA1I", "CALM1", "HRH1", "HTR1A", "HTR2A", "KCNA10", "KCNH2"] | 0.0010296 | 0.017378 |
| Piperlongumine | ["Glutathione transferase inhibitor"] | String[] | 0.0747485 | 0.0144167 |
| Podophyllotoxin | ["Microtubule inhibitor", "Tubulin inhibitor"] | ["IGF1R", "CASP3", "TOP2A", "TUBA4A", "TUBB"] | 0.0010296 | 0 |
| Prazosin hydrochloride | ["Adrenergic receptor antagonist"] | ["ADRA1A", "ADRA1B", "ADRA1D", "ADRA2A", "ADRA2B", "ADRA2C", "CDK1", "KCNH2", "KCNH6", "KCNH7"] | 0 | 0.000328 |
| Progesterone | | ["PGR", "CYP17A1", "NR3C2", "CATSPER1", "CATSPER2", "CATSPER3", "CATSPER4", "CYP2C19", "ESR1", "OPRK1", "TRPC5"] | 0 | 0 |
| Proguanil | ["Dihydrofolate reductase inhibitor"] | ["CYP2C19", "DHFR"] | 0.0016163 | 0 |
| Promazine hydrochloride | ["Dopamine receptor antagonist"] | ["CHRM5", "DRD2", "ADRA1A", "ADRA1B", "ADRA1D", "CHRM1", "CHRM2", "CHRM3", "CHRM4", "DRD1", "DRD3", "DRD4", "HRH1", "HTR2A", "HTR2C"] | 0.024341 | 0 |
| Propafenone hydrochloride | ["Antiarrhythmic"] | ["KCNH2", "SCN5A", "ADRB1", "ADRB2", "KCNA5", "KCNK2", "KCNK3"] | 0 | 0 |
| Propionylpromazine hydrochloride | | | 0 | 0 |
| Protriptyline hydrochloride | ["Tricyclic antidepressant"] | ["SLC6A2", "SLC6A4"] | 0 | 0 |
| Psoralidin | | | 0 | 0 |
| Pyridostatin trifluoroacetate salt | | | 0 | 0.1911159 |
| Quinacrine dihydrochloride | | | | |
| Quinidine sulfate | | | | 0 |
| RN-9893 | | | 0.2081328 | 0.1041704 |
| RU-SKI 43 maleate | | | 0 | 0 |
| Rabeprazole sodium | ["ATPase inhibitor", "Gastrin inhibitor"] | ["ATP4A", "CYP2C19"] | 0.0919194 | 0 |

| CompoundName | MOA | Targets | RMPV750 | RMPV1500 |
|---|---|---|---|---|
| Raloxifene hydrochloride | ["Estrogen receptor antagonist", "Selective estrogen receptor modulator (SERM)"] | ["ESR1", "ESR2", "ACVRL1", "ENG"] | 0 | 0 |
| Ranolazine dihydrochloride | ["Sodium channel blocker"] | ["SCN9A", "SCN10A", "SCN5A", "SLC22A2"] | 0.0022114 | 0.0208727 |
| Reserpine | ["Vesicular monoamine transporter inhibitor"] | ["SLC18A2", "SLC18A1"] | 0 | 0 |
| Ro 11-1464 | | | 0.0028112 | 0 |
| Ro 90-7501 | ["Beta amyloid inhibitor"] | ["APP"] | 0 | 0.0012478 |
| Roscovitine | ["CDK inhibitor"] | ["CDK2", "CDK9", "CDK7", "CDK1", "CDK5"] | 0 | 0 |
| Rotenone | | ["MT-ND1"] | 0.0003682 | 0 |
| Ruthenium red | | | 0.0626678 | 0 |
| S-(+)-Fluoxetine hydrochloride | | | 0 | 0.2889133 |
| S-Methylisothiourea hemisulfate | | | 0.051782 | 0.218206 |
| SB 202190 | ["p38 MAPK inhibitor"] | ["MAPK14", "AKT1", "ALOX5", "CHEK1", "GSK3B", "LCK", "MAPK1", "MAPK11", "MAPK12", "MAPK8", "PRKCA", "ROCK1", "RPS6KB1", "SGK1"] | 0 | 0 |
| SB 415286 | ["Glycogen synthase kinase inhibitor"] | ["GSK3B", "GSK3A", "RPS6KB1"] | 0 | 0.0012478 |
| SB743921 hydrochloride | | | 0 | 0 |
| SID 3712249 | | | 0.002513 | 0 |
| SKF 83959 hydrobromide | | | 0.0456887 | 0 |
| SMER28 | | | 0 | 0 |
| SP600125 | | | 0 | 0 |
| SR 59230A oxalate | ["Adrenergic receptor antagonist"] | ["ADRB3", "ADRB1", "ADRB2"] | 0.0106818 | 0 |
| SR9243 | | | 0.0694414 | 0.0012478 |
| SU 5416 | | | 0.0212056 | 0.3580667 |
| SU1498 | | | 0 | 0 |
| Sanguinarine chloride | | | | |
| Sertaconazole nitrate | ["Sterol demethylase inhibitor"] | String[] | 0.0119337 | 0 |
| Stattic | | | | |
| Sunitinib | ["FLT3 inhibitor", "KIT inhibitor", "PDGFR receptor inhibitor", "RET tyrosine kinase inhibitor", "VEGFR inhibitor"] | ["FLT3", "KDR", "KIT", "FLT4", "FLT1", "PDGFRA", "PDGFRB", "RET", "CSF1R", "FGFR1"] | 0 | 0.3831555 |
| Supercinnamaldehyde | | | 0 | 0.0144167 |
| Suprafenacine | | | 0 | 0 |
| T0070907 | | | 0 | 0 |
| TBBz | | | 0.0010296 | 0.0026091 |
| TIC10 angular | | | 0 | 0 |
| Tacrine | ["Acetylcholinesterase inhibitor", "Acetylcholine release stimulant", "Butyrylcholinesterase inhibitor", "Potassium channel antagonist"] | ["ACHE", "BCHE"] | 0 | 0 |
| Tamoxifen citrate | | | | |

| CompoundName | MOA | Targets | RMPV750 | RMPV1500 |
|---|---|---|---|---|
| Taurine | | | 0 | 0.4691108 |
| Testosterone | ["Androgen receptor agonist"] | ["AR", "CYP19A1", "CYP2C19", "CYP2C8", "CYP3A5"] | 0.0125635 | 0.0554207 |
| Tetraethylthiuram disulfide | | | | |
| Thiabendazole | | | 0.010286 | 0.0012478 |
| Thiocolchicine | | | 0 | 0 |
| Tirapazamine | | | 0.0007038 | 0.0020819 |
| Tizanidine hydrochloride | ["Adrenergic receptor agonist"] | ["ADRA2A", "ADRA2B", "ADRA2C", "CYP1A2", "NISCH"] | 0 | 0 |
| Tolazoline | ["Adrenergic receptor antagonist"] | ["ADRA2A", "ADRA2B", "ADRA2C", "ADRA1A", "HRH1", "HRH2"] | 0.0338009 | 0.1041704 |
| Topotecan hydrochloride hydrate | ["Topoisomerase inhibitor"] | ["TOP1", "TOP1MT"] | 0 | 0 |
| Torin2 | | | 0 | 0 |
| Trifluperidol hydrochloride | | | 0.0007038 | 0 |
| Triflupromazine hydrochloride | ["Dopamine receptor antagonist"] | ["HTR2B", "CHRM1", "CHRM2", "CHRNA7", "DRD1", "DRD2"] | 0 | 0 |
| Trihexyphenidyl | | ["CHRM1", "CHRM2", "CHRM3", "CHRM4", "CHRM5"] | 0.0467328 | 0.1044492 |
| Trilostane | | | 0 | 0 |
| Trimipramine maleate | ["Norepinephrine reuptake inhibitor", "Tricyclic antidepressant"] | ["SLC6A2", "SLC6A4", "SLC6A3", "ADRA1A", "ADRA1B", "ADRA2A", "ADRA2B", "ADRB1", "ADRB2", "ADRB3", "CHRM1", "CHRM2", "CHRM3", "CHRM4", "CHRM5", "DRD1", "DRD2", "DRD5", "HRH1", "HTR1A", "HTR1D", "HTR2A", "HTR2C", "HTR3A"] | 0.0216222 | 0 |
| Tyrphostin AG 879 | | | 0 | 0 |
| U-101958 maleate | | | 0.3587426 | 0.1732954 |
| U-62066 | | | 0.562672 | 0.0015266 |
| U0126 | | ["JAK2", "MAP2K1", "MAP2K2", "MAP3K1", "MAP3K2"] | 0 | 0 |
| UNC0379 trifluoroacetate salt | | | 0.0013317 | 0 |
| Vinblastine | ["Microtubule inhibitor", "Tubulin inhibitor"] | ["TUBB", "JUN", "TUBA1A", "TUBD1", "TUBE1", "TUBG1"] | 0 | 0.1859498 |
| Vincristine sulfate | | ["TUBB", "TUBA4A"] | 0.0003682 | 0 |
| Vorinostat | | | | |
| WIN 62,577 | | | 0.0100583 | 0.0020819 |
| WZ4003 | | | 0 | 0 |
| Wiskostatin | ["Neural Wiskott-Aldrich syndrome protein inhibitor"] | ["WAS", "WASL"] | 0 | 0 |
| XL388 | | | 0 | 0 |
| Y-27632 dihydrochloride | ["Rho associated kinase inhibitor"] | ["ROCK1", "ROCK2", "LRRK2", "PKIA", "PKN2", "PRKACA", "PRKCE"] | 0.0267891 | 0.0056832 |
| Yoda1 | | | 0 | 0.0443088 |

| CompoundName | MOA | Targets | RMPV750 | RMPV1500 |
|---|---|---|---|---|
| alpha-Lobeline hydrochloride | | | 0.0106818 | 0.0347489 |
| beta-Lapachone | | | | |

**Supp. Table 1: Compound list in chemical HCS experiment.** Description of the name of the screened compounds (CompoundName), their mechanisms of action (MOA) and genetic targets (Targets) as well as their robust morphological perturbation value (FDR-corrected p-value for a comparison to matching DMSO controls) computed on the plate seeded with 750 cells (RMPV750) or 1500 cells (RMPV1500).
